# Leech Speciation and Detection of Hirudin Gene among the Leech Population in Asejire Oyo State, Nigeria

**DOI:** 10.1101/2025.06.02.657427

**Authors:** T.H. Olajide, A.O. Ajileye, O.J. Bamikole, M.-D.B. Olufeagba, B.A. Adedeji, S.A. Ademola, O.K. Amodu

## Abstract

Leeches are segmented, parasitic and sanguivorous worms of the family Hirudinae and phylum Annelida. Various ailments have been treated with them since ancient times. Recent evidence has established the presence of over 100 medicinally useful bioactive substances in their saliva. Based on the therapeutic properties of the substances, they have been used in different areas of medical care and most prominently in Plastic and Reconstructive Surgery. Wounds and injuries account for more than a third of hospital visits and as much as 9% annual mortality in Africa. Nigeria with her abundant leech population however still accounts for the bulk of the population living with conditions treatable with leech therapy. Hence, this study aimed at confirming the presence of Hirudin and identifying the species of Leeches collected from Asejire, Ibadan.

Fifty Leeches were collected from Asejire, Ibadan, Oyo State. Ethical approval (AD 13/479/4346^B^) was obtained from the Oyo State ethical board, Identification was morphologically done using the Borda and Siddall classification criteria. DNA extraction and amplification for Hirudin gene was similarly done using PCR. The mitochondrial cytochrome oxidase subunit 1 (COI) and nuclear internal transcribed spacer-2 (ITS2) genes were sequenced using Sanger sequencing. Multiple Sequence Alignments were done using the MUSCLE alignment method and phylogenetic trees were constructed via MEGA-x software using the Kimura-2 parameter model and Maximum Likelihood method.

The ITS2, COI and Hirudin genes were characterized at 517bp, 658bp and 226bp respectively. A BLAST search showed the COI gene in our local Leech species is most closely related to the *Aliolimnatis michaelsenis* with 94.50%, nucleotide similarity, and the ITS2 is similar to *Poecilobdella (Hirudinaria) manillensis* with a 97.18% nucleotide similarity.

This study confirms the presence of Hirudin in the Leech species found in our locality thus their feasibility for use in Leech therapy. Taken together, the phylogenetic inferences showed the Leech Species in our population is a unique previously unidentified species. Further study is required to determine other active compounds in our unique species and how it can be harnessed for medical use.

## INTRODUCTION

Leeches are segmented, parasitic and sanguivorous worms, belonging to the family *Hirudinae* and phylum *Annelida* (Buchsbaum *et al*, 1987). The family *Hirudinidae* (*Arhynchobdellida, Hirudiniformes*) is comprised of mainly blood-sucking (*sanguivorous*) freshwater leeches, or medicinal leeches, although four terrestrial species are known (Elliott and Kutschera 2011).

Leech therapy is the process of using leeches to improve blood circulation to a wound site and preventing tissue death (Alaama et al., 2014) has been used as a healing technique for centuries to treat diseases till the twentieth century when it became unpopular in the medical community. Recent advances in scientific research globally over the last 30 years have witnessed the increased attention and resources invested in leech therapy and leeching research. The medicinal leeches are among the invertebrates with the best studied biology and physiology because of their wide application in medicine (Todorov et al., 2016; Sig et al., 2017).

Leeching is becoming more popular in medicine, notably in plastic and reconstructive surgery owing to the anticoagulant, thrombolytic and vasodilant properties of their saliva (Abdullah *et al*, 2012), the leech saliva contains over 100 bioactive compounds; Hirudin, Hyaluronidase, Destabilase, Apyrase, Eglin, Calin, Guamerin among others which have been found to have anticoagulant, vasodilatant, anesthetic, thrombolytic, antibiotic, analgesic, anti-metastatic and anti-inflammatory properties hence, their medicinal uses (Zaidi et al., 2011). Leeches accomplish this by sucking clotted blood from sensitive regions like beneath a flap of skin or on a finger or toe (Singh and Rajoria, 2020). This improves blood flow in the small blood vessels and aids in the prevention of tissue death. Leech therapy has been employed in the treatment of a wide range of other diseases such as skin pathologies, nervous system abnormalities, urinary and reproductive system problems, and inflammation and dental problems (Ojo et al., 2010).

In 2004, the United States Food and Drug Administration (FDA) granted market approval for medicinal leech therapy as “An adjunct to the healing of graft tissue when problems of venous congestion may delay healing, or to overcome problems of venous congestion by creating prolonged bleeding” (Koeppen *et al*, 2019).

Hirudin, the most popular and most powerful anticoagulant compound isolated from leeches has been used as an anticoagulant in many renal diseases that include chronic renal failure, glomerulonephritis, diabetic nephropathy, ulcers and chronic wounds (Glusa, 1998). Wounds and injuries accounting for 30 – 42% of hospital attendance are one of the leading causes of hospital visits in Africa, it is implicated in about 9% of global annual deaths (Norman et al, 2006; Builders and Momodu, 2017). Several studies conducted by the World Health Organization has found an increase in individuals seeking alternative wound treatment methods as to the available conventional treatment methods most especially in Low and Middle-Income Countries like Nigeria (WHO, 2019). Leech therapy is a good and safe example of alternative treatment in wound management.

Nigeria, although blessed with an abundance of freshwater bodies and damp sites that accommodate several organisms including Leeches and plagued with medical conditions in which leeching has been proven to be therapeutic, is yet to tap into the huge potentials of leeching. Thus, we sought to find out if the Leech species present in our locale are medicinal and ascertain the suitability of these Leech species for use in Leech therapy. This would consequently bring about a reduction in the cost of wound treatment, speed up wound healing time, prevent the need for subsequent procedures (in the case of plastic surgeries), prevent untimely skin or body disfiguration and avoid gradual progression of chronic wounds to keloids and cancer. This study aimed to genetically identify Leech species and determine the presence of Hirudin in the identified species.

## METHODS

Fifty (50) Leeches were randomly collected from freshwater habitats and damp surfaces in Asejire Lake is located in Southwestern Nigeria. The leeches were maintained in well-aerated plastic containers filled with un-chlorinated tap water. Water in the container was regularly changed at two-day intervals and kept at room temperature (25°C). morphological and Taxonomic identification of the collected leeches were confirmed by visual inspection (body coloration pattern) using Nikon SMZ-10 binocular stereo microscope according to Borda and Siddall (2004);Elliot and Kutschera (2011). Genomic DNA was isolated using the Quick DNA™ Miniprep Plus Kit (Zymo Research). Concentration and purity of DNA samples were ascertained to determine their yield using NanodropOne_AZY1810399. PCR amplification using universal primers 5’-GCATCGATGAAGAACGCAGC-3’ and 5’-TCCTCCGCTTATTGATATGC-3’, 5’-GGTCAACAAATCATAAAGATATTGG-3’ and 5’-TAAACTTCAGGGTGACCAAAAAATCA-3’. was carried out according to the conditions of Trontelj and Utevsky (2005). Hirudin detection was carried out by amplification using (5′-AGTGCATATTGGGTTCTAATGGA-3′) and reverse (5′TGGTAAATAGCTGAATATGATTGAGAG-3′) primers following the protocol of Al-badran and Al-fadal (2017). Amplicons were viewed on 2% agarose. Automated nucleotide sequencing was performed on an ABI 3130XL Nucleotide Sequence Analyzer via Sanger sequencing. The sequences were edited with Sequence Scanner software, version 1.0 (Applied Biosystems, Foster City, CA) and deposited in the GenBank. The nucleotide sequences comparison was done using Basic Local Alignment Search Tool (BLAST) (Altschul, 1990) via the National Center of Biotechnology Information. Multiple sequence alignments between the Nigerian Leech species COI and ITS2 gene sequences and sequences retrieved from the GenBank were carried out with Muscle algorithm in BioEdit Sequence Alignment Editor Software (version 7.2.5; Hall, 1999). The phylogenetic relationships between haplotypes of the leeches were reconstructed using the Maximum Likelihood method under the Kimura 2-parameter model (Kimura, 1980). Corresponding sequences were used to root the phylogenetic tree as an outgroup. Confidence in estimated relationship was determined using the bootstrap approach obtained through 1000 replicates with the same model as mentioned above (Felsenstein, 1985). Both bootstrap analysis and phylogeny reconstruction were conducted using MEGA-x (version 10.2.6).

## RESULTS

Upon amplification of the Hirudin gene, bands were present and positioned at 226bp. Bands were clearly visible at 517 base pairs (bp) for the amplification of ITS2 rRNA region while that of the mitochondrial COI region was visible at 658bp.

The three ITS2 sequences compared with sequences in the GenBank database using BLAST showed a 97.18%, 97.92% and 97.26% nucleotide percentage identity each with Accession numbers KX215702.1, JX885697.1, JX885696.1, JX885695.1 and JX885694. The five COI sequences showed nucleotide percentage identities of 94.50%, 94.66%, 94.75%, 94.52% and 94.17% with existing sequences in the GenBank database. Constructed phylogenetic tree showed a difference between our leech specie and globally available species on Genbank.

## DISCUSSION

Leech therapy has been a well-known therapeutic practice throughout the ages for a wide range for diseases and applied as an unorthodox home-remedy by traditional therapists. Today, leeching is being used in the orthodox medical setting with fewer applications, which have been proven and supported by a large number of scientific studies and case reports (Scacheri et al, 1993; Markwardt, 2002; Galtier et al, 2009; Singh, 2010; Zaidi et al, 2011; Abdullah et al, 2012; Dwivedi, 2012; Okka, 2013; Das, 2014; Mumcuoglu, 2014; Rahul et al, 2014; Hildebrandt, 2011; Sig et al, 2017; Kvist et al, 2020). Several nuclear gene (ITS1, ITS2, 18S rRNA and 28S rRNA) and mitochondrial gene (COI, 12S rDNA) amplification have been useful in the identification of leech species and phylogenetic analysis (Trontelj and Utevsky, 2005; Zivic *et al*, 2015; Won *et al*, 2014; Müller *et al*, 2015; Bilal *et al*, 2017).

This study was carried out to investigate the type of leeches found in Asejire community, Ibadan, Nigeria and their feasibility for use in Leech therapy. Hirudin gene present in leeches was successfully characterized at 226bp in this study. This showed that the Leech species in our locality possess the medicinal hirudin properties, hence can be termed medicinal Leeches and thus used in the achievement of Leech therapy. Our study focused on the amplification of a nuclear gene and a mitochondrial gene to genotype leech species (ITS2 and COI genes). Cladistic analysis employed 517bp of nuclear ITS2 ribosomal DNA and 658bp of mitochondrial cytochrome C oxidase subunit I, in addition to morphological data. Apakupakul *et al*, (1999) suggested that the use of two molecular data sets, one nuclear gene and one mitochondrial gene as well as morphological data, combined historical information evolving under a variety of different constraints and therefore was less susceptible to the biases that could confound the use of only one type of data. Our study suggests that the nuclear ITS2 gene yields a meaningful historical signal for determining higher level relationships. The more rapidly evolving COI gene was informative for recent or local areas of the evolutionary hypothesis, such as within-family relationships.

Sequence alignment and estimation of genetic difference are crucial steps in molecular evolutionary studies. Sequence analysis of the leech species in our locality, using the BLAST algorithm for COI gene, showed 94.50%, 94.66%, 94.75%, 94.52% and 94.17% nucleotide identity with *Aliolimnatis michaelseni*. This corroborates claims by Omalu *et al* (2015), Omalu *et al* (2016) and Mgbemena *et al*, (2020) that the Leech samples collected from the North Central geographical zone of Nigeria are *A. michaelseni*. However, phylogenetic relationships between Leech species retrieved from the GenBank and the Leech species in our locale showed the possibility of a new species. For ITS2 gene analysis of our local Leeches, the three sequences had 97.18%, 97.92% and 97.26% similarity to *Poecilobdella* (*Hirudinaria) manillensis* isolate (KX215702.1). *H. manillensis* is among the commonly used medicinal leeches (Thakur *et al*, 2017) and thus adds credence to the possibility of Leech therapy with the Leech species found in our locale. The three sequences of the ITS gotten from this population are presently in GenBank database and assigned the accession numbers ON209372, ON209373, and ON209374.

An important outcome of the molecular analysis of our work revealed that our specie is suitable for medicinal leech therapy. This is in contrast to the age-long belief that all medicinal Leeches are *Hirudo medicinalis*. This outcome corroborates several studies that have shown other leech species outside the class Hirudinae have medicinal capabilities (Siddall and Burreson, 1998; Apakupakul *et al*, 1999; Utevsky and Trontelj, 2005; Siddall *et al*, 2007; Phillips, 2012). Todorov *et al*., (2016) asserts that while the widespread application of Leeches in medicine is probably the main reason for it being among the invertebrates with the most studied biology and physiology, the same cannot be said of Leeches with regard to their taxonomy which has been substantially revised in recent years through modern molecular and cytogenetic methods. This has necessitated the need for a review of the systematics of the Leech taxa.

## CONCLUSION

This study confirms the presence of Hirudin, a major bioactive substance in Leech therapy, in the Leech species found in our locale and as such are qualified to be called Medicinal Leeches. This makes the prospect of achieving leech therapy from our local Leech population highly feasible. The study has also shown the possibility of a new species whose sequences have been deposited at the GenBank database. There is no record of any species sequence from Nigeria therefore, the submission of our sequences to the GenBank database would add to literature as well as serve as a future reference material in the knowledge of Leech species from Nigeria.

---

**Table 1:** The Common Composition of Leech Saliva.

| Chemical constituent | Mode of action |
| --- | --- |
| Hirudin | Inhibits blood coagulation by binding to thrombin. |
| Calin | Inhibits blood coagulation by blocking the binding of Von Willebrand factor to collagen, inhibit collagen mediated platelet aggregation |
| Destabilase | It has glycosidase activity and its monomerizing activity dissolves fibrins. It also has antimicrobial activity |
| Hirustatin | Inhibits kallikrein, trypsin, chymotrypsin and neuropholic cathepsin G and also accelerates reperfusion and prevents reocclusion in a canine model of femoral arterial thrombosis. |
| Bdellins | Anti-inflammatory inhibits plasmin, trypsin and acrosin. |
| Hyaluronidase | It reduces the viscosity and renders the tissues more readily permeable to injected fluids, increasing the speed of absorption. This promotes resorption of excess fluids and extravasated blood. |
| Tryptase Inhibitor | Inhibits proteolytic enzymes of host's mast cells. |
| Eglins | Anti-inflammatory, inhibits the activity of alpha chymotrypsin chymase,subtilisin,elastase,cathepsin G. |
| Factor Xa inhibitor | Inhibits the activity of coagulation factor Xa by forming equimolar complexes. |
| Carboxypeptidase | Increase the inflow of blood at the bite site of inhibitors |
| Acetylcholine | Increase blood flow by dilating constricted vessels |

**Figure 1:**
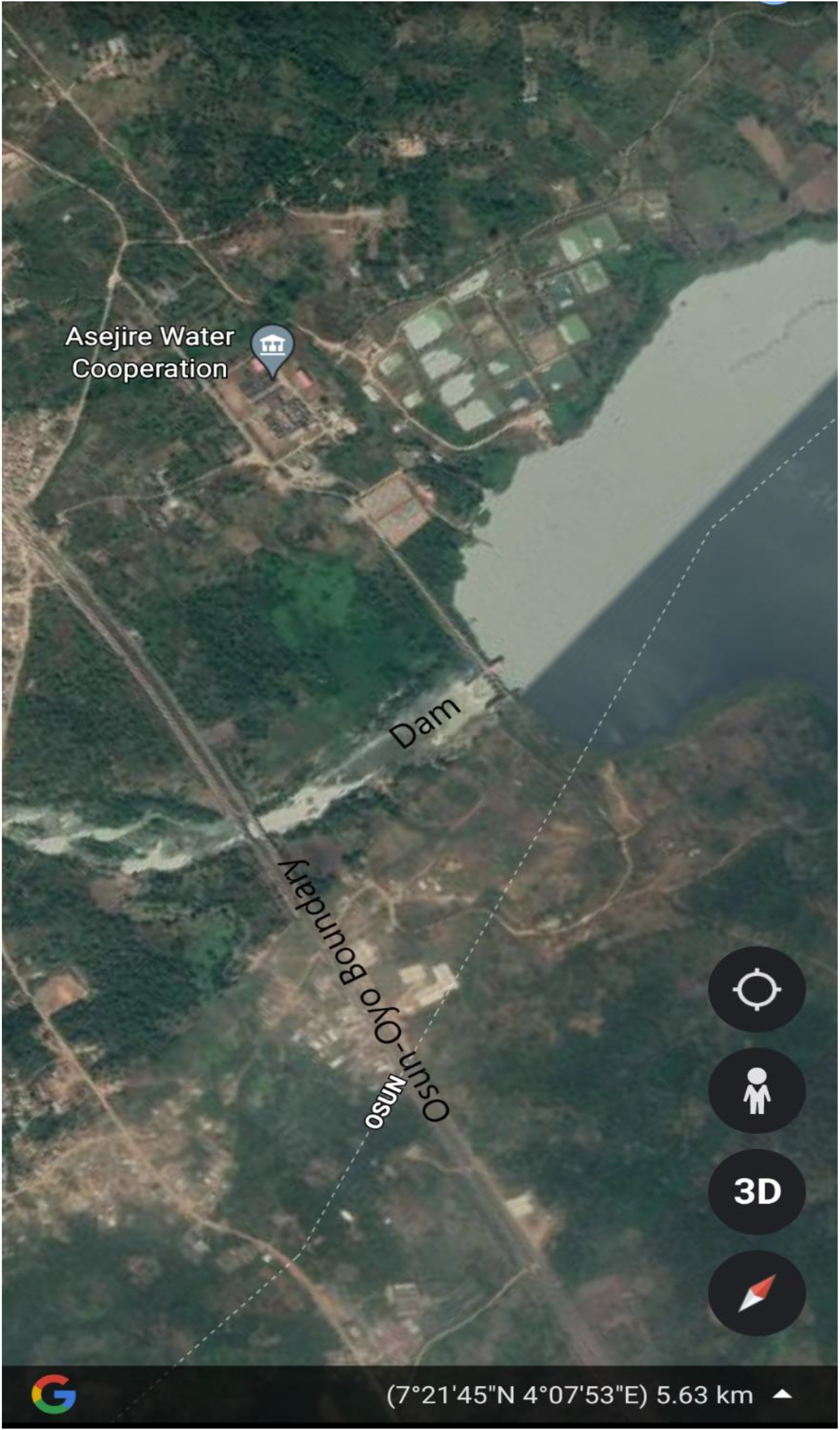
Map of Osun River Basin showing Asejire reservoir (Google Earth)

**Figure 2a:**
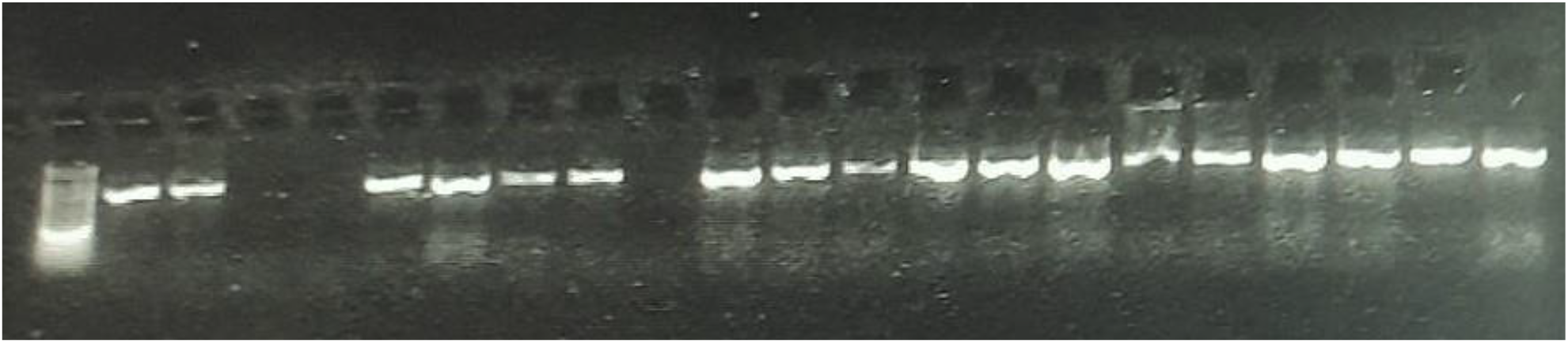
Gel Electrophoresis showing the amplification of ITS2 gene at 517bp from DNA samples extracted from Leech species.

**Figure 2b:**
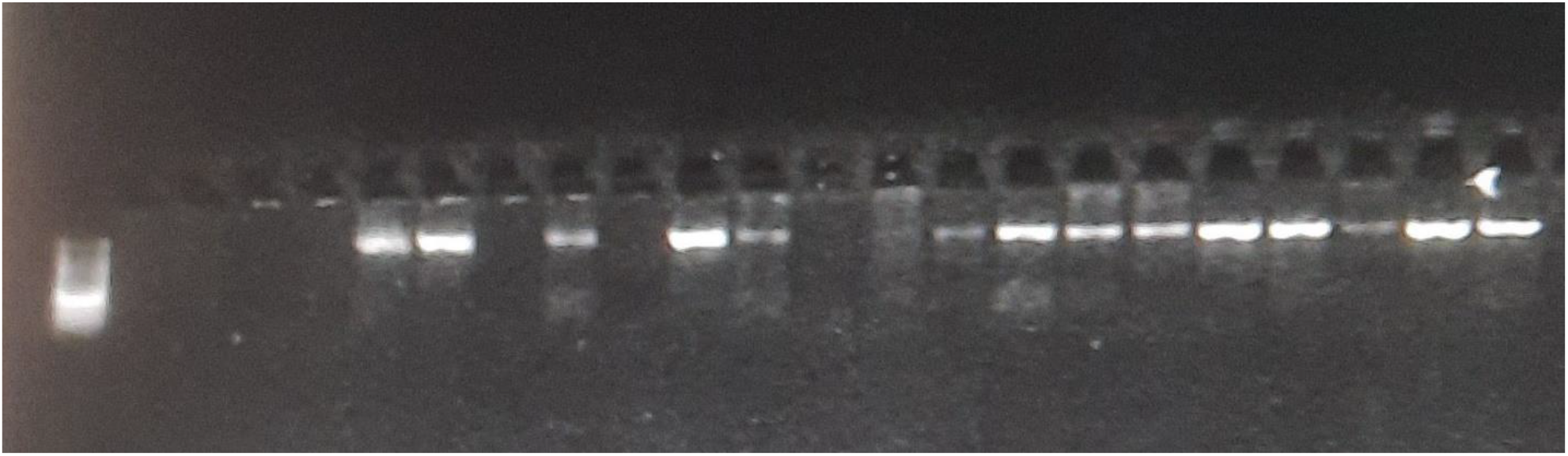
Gel Electrophoresis showing the amplification of COI gene at 658bp from DNA samples extracted from Leech species.

**Figure 2c:**
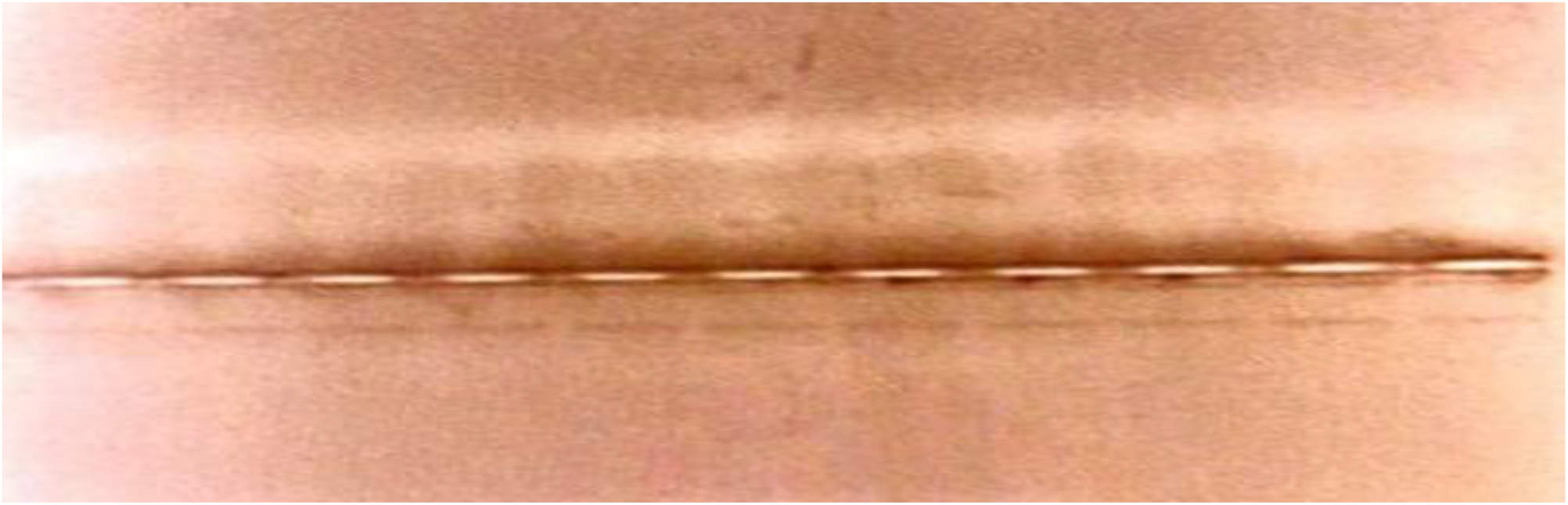
Gel Electrophoresis showing the amplification of Hirudin gene at 226bp from DNA samples extracted from Leech species.

**Figure 4.5a–c:**
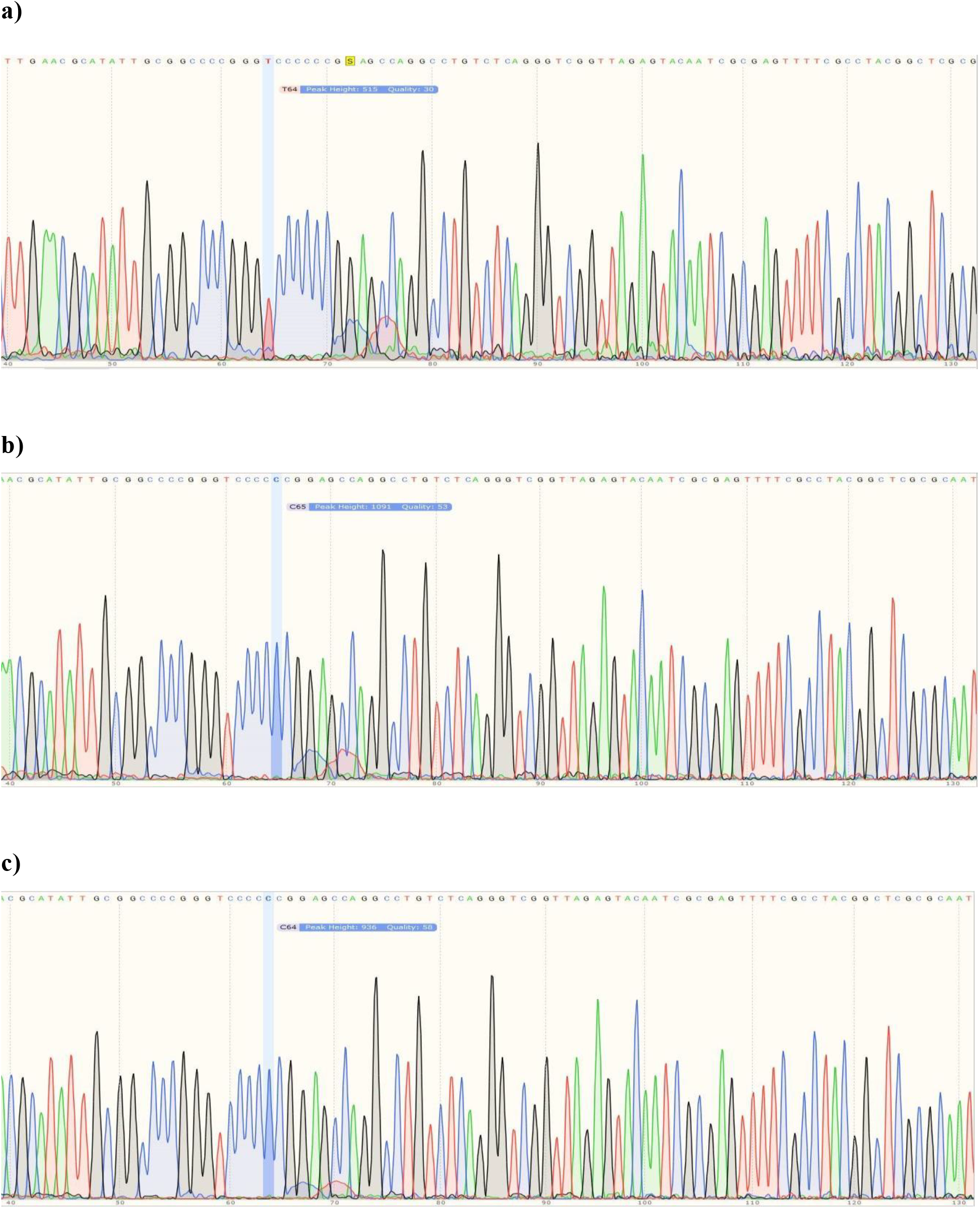
Sanger DNA sequencing of our Leech Species showing their chromatograms (for ITS2 gene)

**Figure 4.5d–f:**
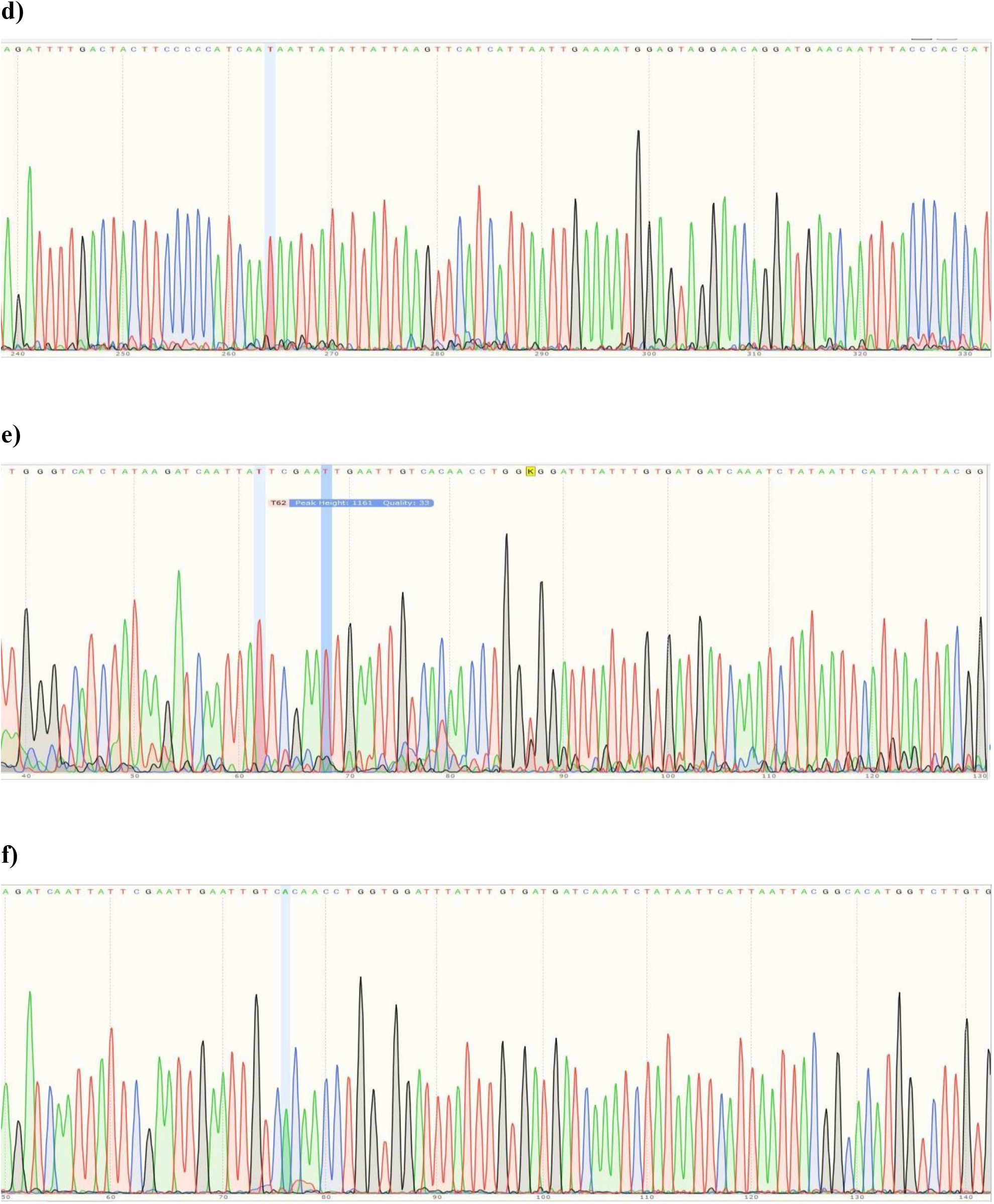
Sanger DNA sequencing of our Leech Species showing their chromatograms (for COI gene)

**Figure 3a – h:**
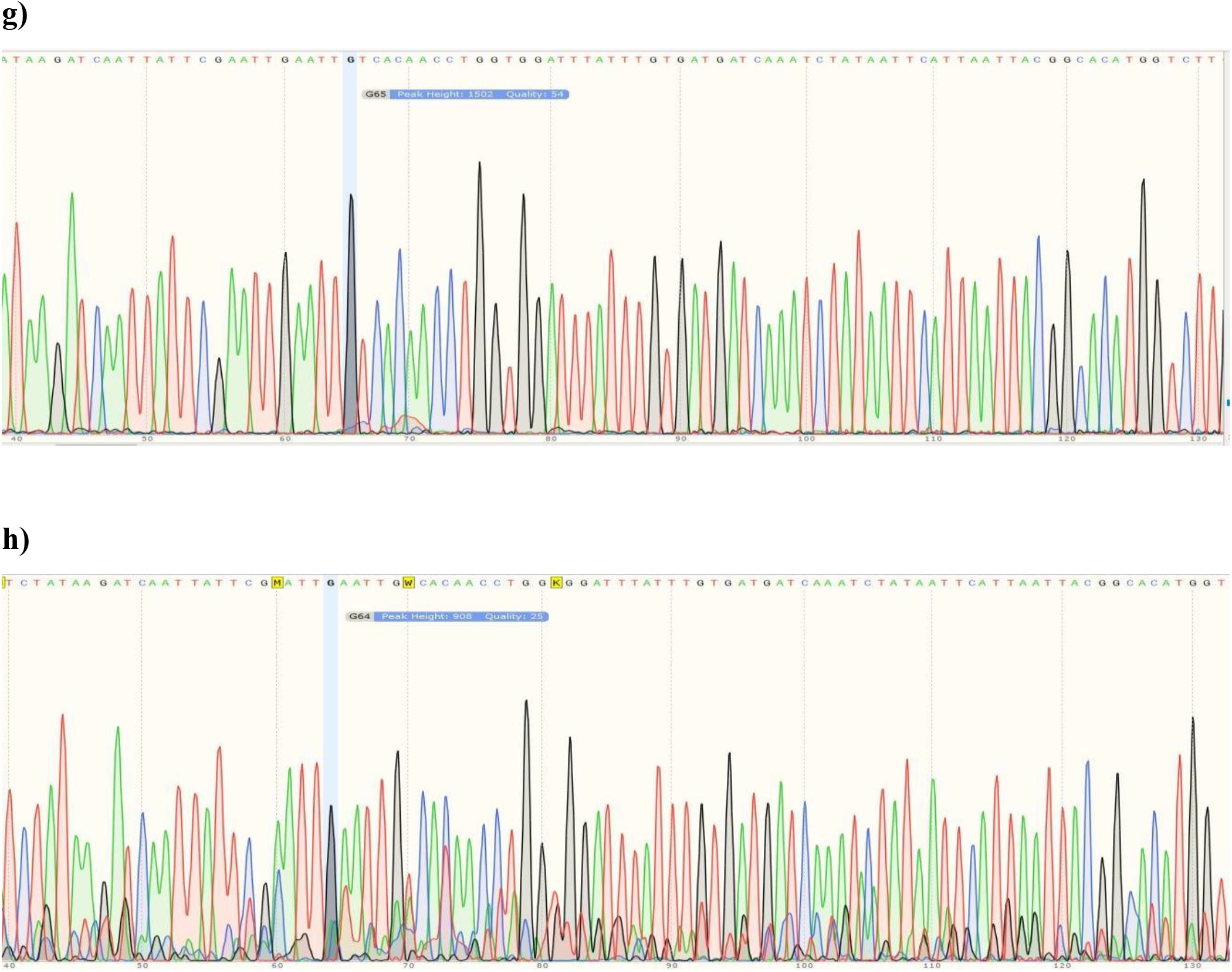
Sanger DNA sequencing of our Leech Species showing their chromatograms (for COI gene)

**Figure 4a – c:**
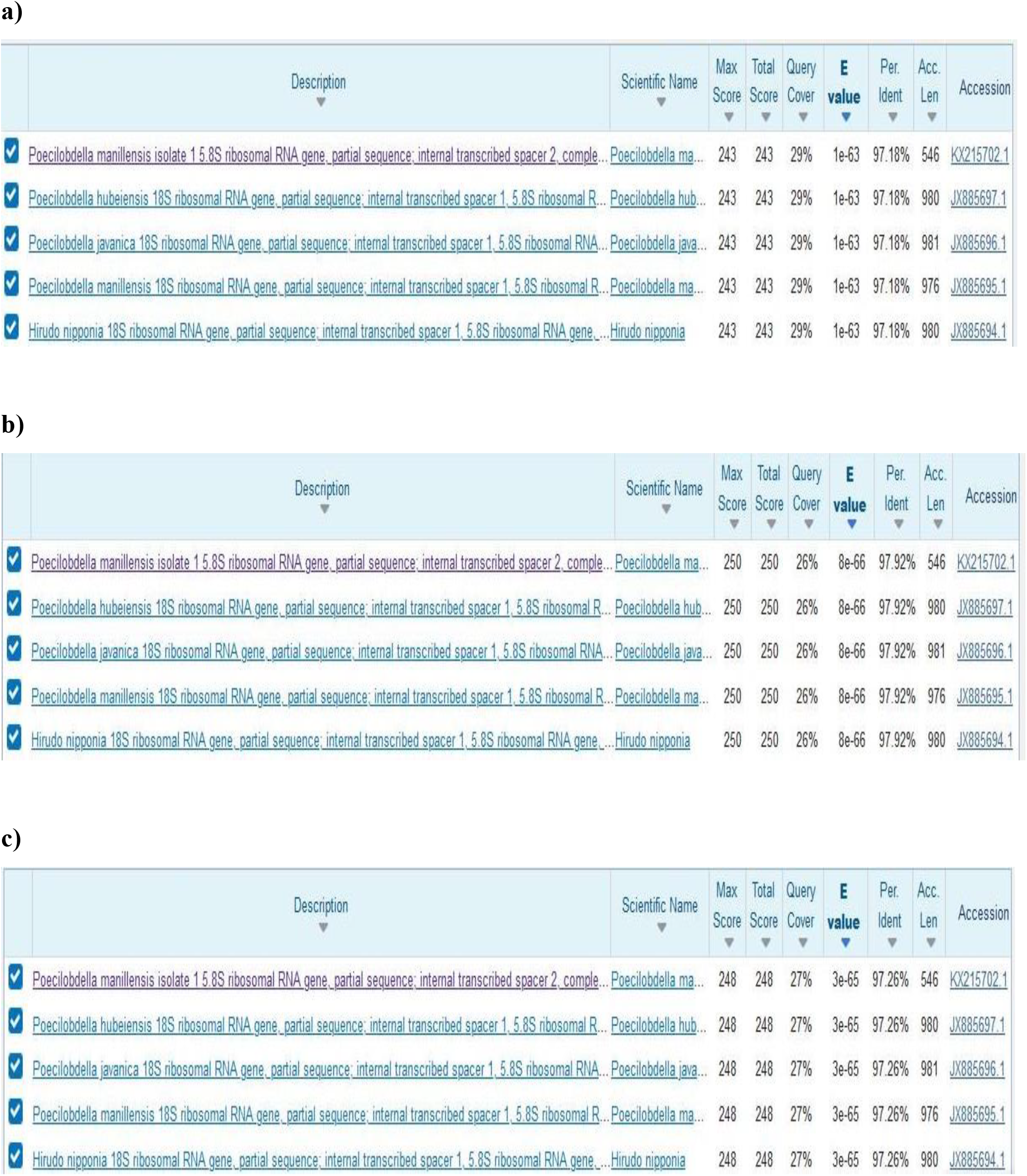
BLAST search between the three ITS2 DNA sequences of our Leech species and Existing Sequences in the Genbank showing a 97.18%, 97.92% and 97.26% identity respectively.

**Figure 5a – e:**
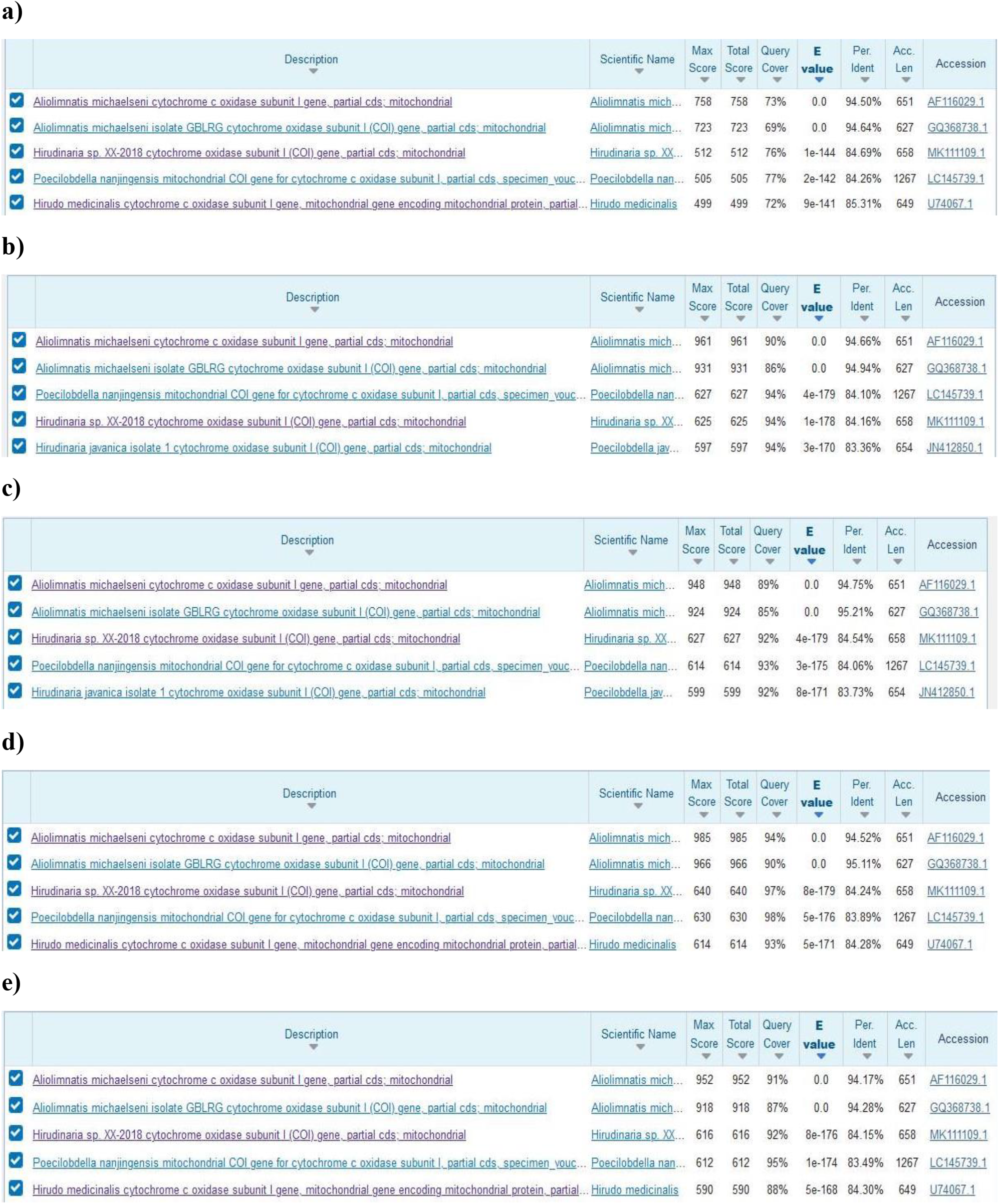
BLAST search between the five COI DNA sequences of our Leech species and Existing Sequences in the Genbank showing a 94.50%, 94.66%, 94.75%, 94.52% and 94.17% highest sequence identity respectively.

**Figure 6a:**
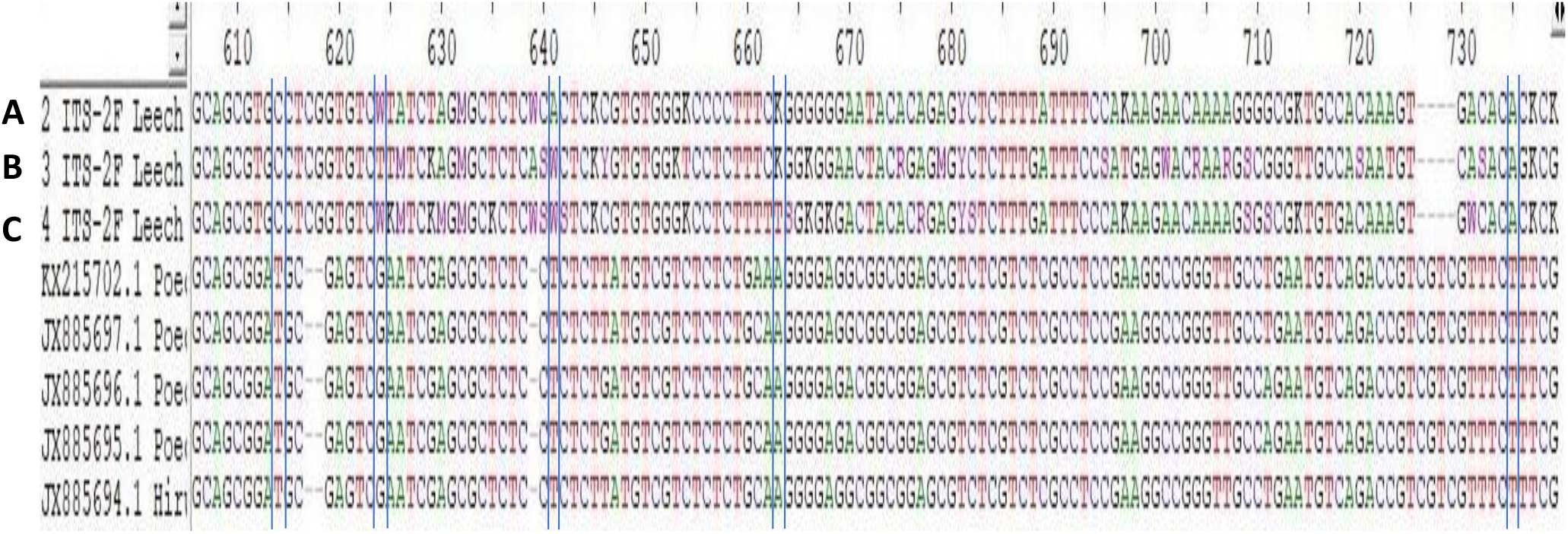
Multiple Sequence alignment of ITS2 Leech DNA sequences and closely related sequences retrieved from GenBank showing several positions of nucleotide mismatch (such as sequences in between the blue lines). Letters A, B and C are sequences from this study

**Figure 6b:**
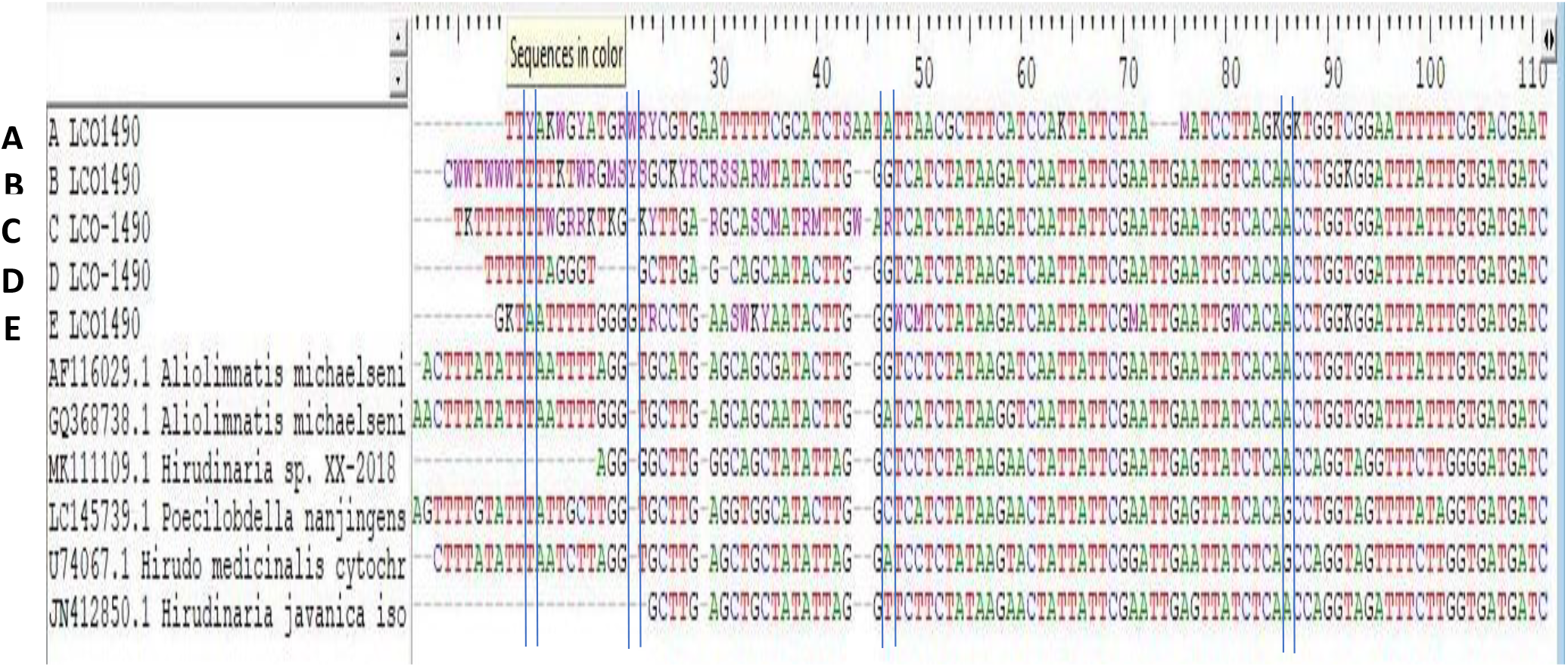
Multiple Sequence alignment of COI Leech DNA sequences and closely related sequences retrieved from GenBank showing several positions of nucleotide mismatch (such as sequences in between the blue lines). Letter A, B, C, D and E are sequences from this study

**Figure 7a:**
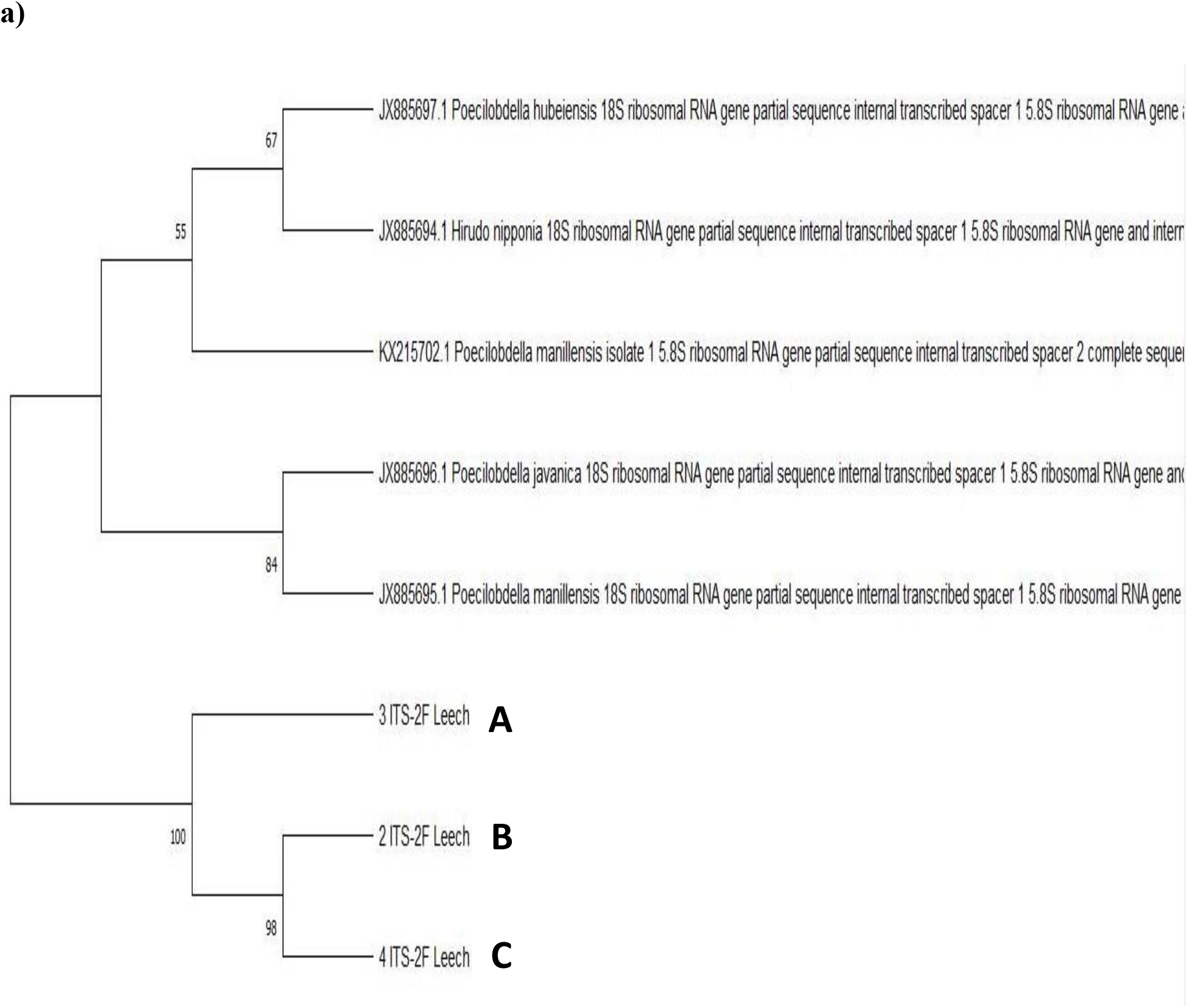
Phylogenetic Tree showing the evolutionary relationship between our Leech species sequences (ITS2) and similar species sequences retrieved from the GenBank via Maximum Likelihood Method implemented in Muscle algorithm with Mega-x. Letters A, B and C are our sequences identified in this study.

**Figure 7b:**
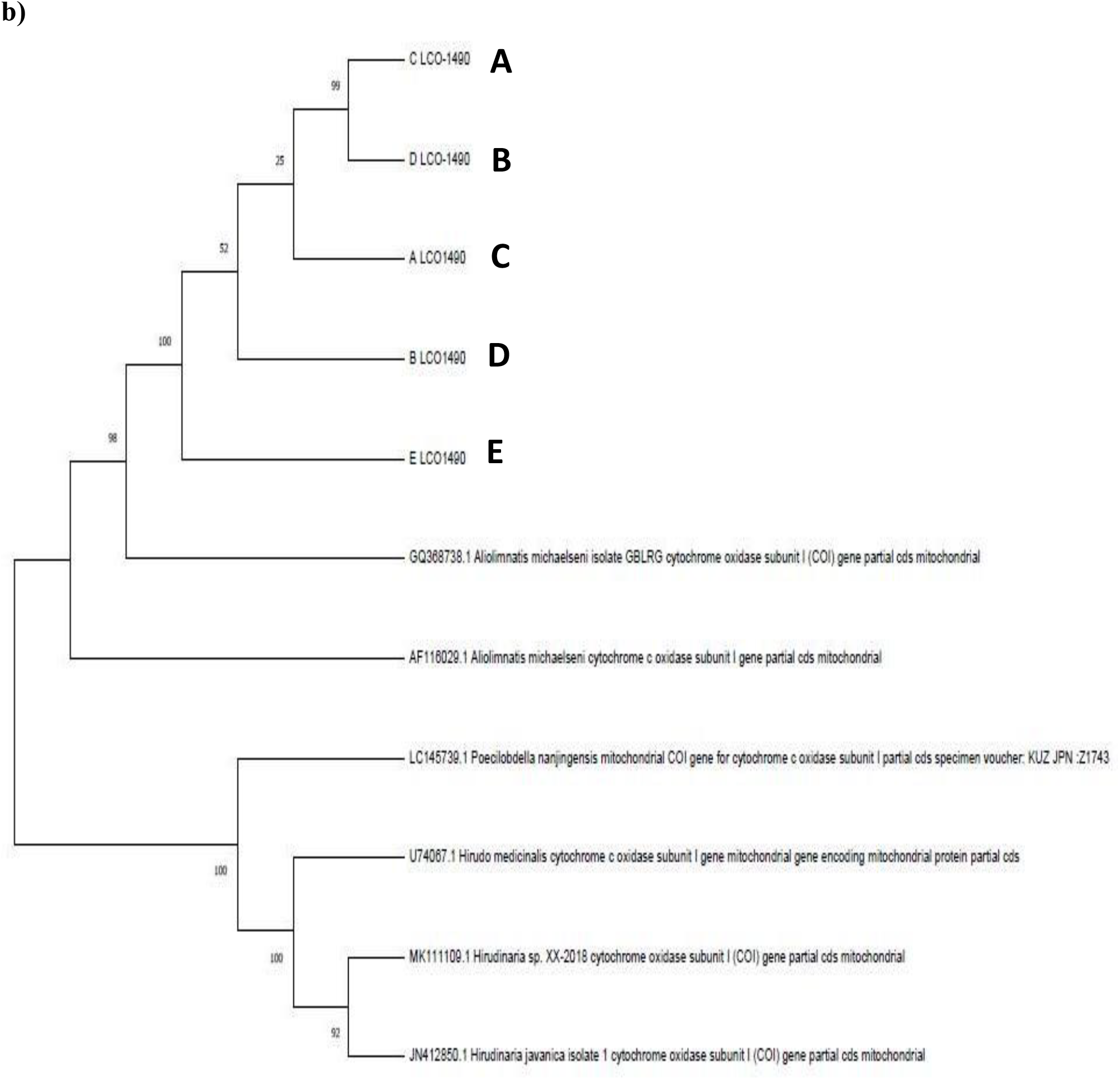
Phylogenetic Tree showing the evolutionary relationship between our Leech species (COI) sequences and similar species sequences retrieved from the GenBank via Maximum Likelihood Method implemented in Muscle algorithm with Mega-x. Letters A, B, C, D and E are our sequences identified in this study.

